## Supplementary information for "Fasciclin 2 functions as an expression-level switch on EGFR to control organ shape and size in *Drosophila*"

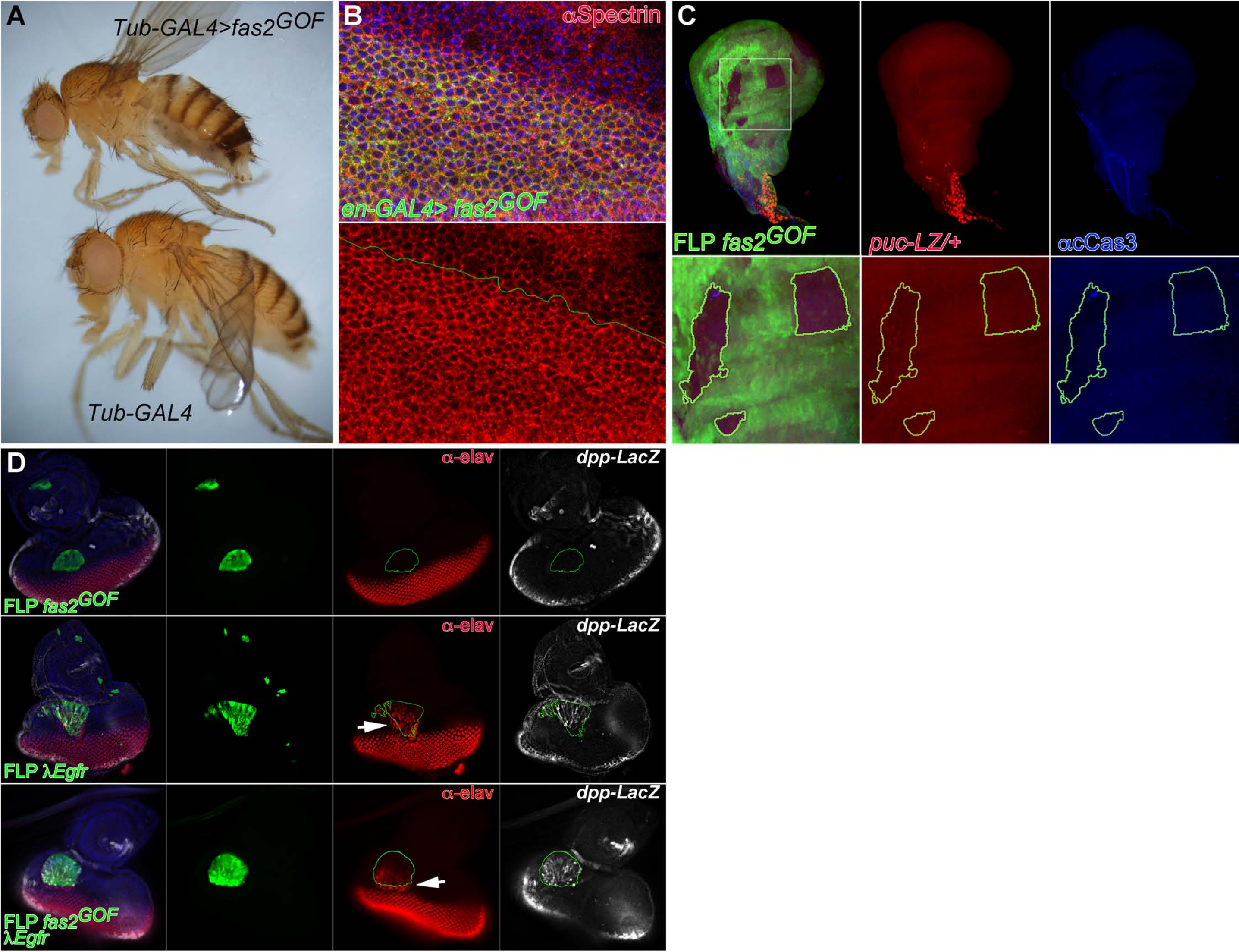

A
Tub-GAL4>fas2GOF
Tub-GAL4
B
αSpectrin
en-GAL4> fas2GOF
C
FLP fas2GOF
puc-LZ/+
αcCas3
D
α-elav
dpp-LacZ
FLP fas2GOF
FLP λEgfr
FLP fas2GOF λEgfr

**S1 Fig. Analysis of the Fas2 GOF condition in adult flies and FLP-OUT clones**

(A) Over-expression of Fas2$^{TRM}$ (*UAS-fas2*$^{TRM}$) under the control of the *Tub-GAL4*/+ driver (*Tub-GAL4>fas2*$^{GOF}$) caused a general reduction of body size. Compare the two females, with over-expression of Fas2$^{TRM}$ (top) and a normal CyO sibling (*Tub-GAL4*) control (bottom).

(B) A/P compartment border in an *en-GAL4 UAS-CD8GFP*/*UAS-fas2*$^{TRM}$ *UAS-fas2*$^{GPI}$ pupal wing. The expression of Fas2$^{GPI}$ and Fas2$^{TRM}$ in the Posterior compartment of the wing (labelled with GFP) did not modify the size or shape of the cells.

(C) FLP-OUT *fas2* GOF clones in a *puc-LacZ*/+ genetic background did not show increased JNK activity nor apoptosis. At left, a wing disc with several Fas2 GOF clones (labeled with GFP) which cover most of the disc surface. The lack of ectopic expression from the reporter *puc-LacZ* reveals that the JNK pathway was not de-repressed in these clones. Staining with anti-cleaved Caspase 3 (Blue channel) did not reveal obvious signs of apoptosis in the wing discs either. Lower panels, note the presence of just three cells expressing cleaved Caspase 3 in one of the normal regions engulfed by the Fas2 GOF clones.

(D) The gain of function for EGFR is epistatic to the Fas2 GOF condition. FLP-OUT *y w hs-FLP; ActinFRTy*$^{+}$*FRT-GAL4 UAS-GFP*/*UAS-fas2*$^{TRM}$ *UAS-fas2*$^{GPI}$; *UAS-λEgfr*/*dpp-LacZ* clones (middle panels) display the same phenotype than FLP-OUT *y w hs-FLP; ActinFRTy*$^{+}$*FRT-GAL4 UAS-GFP*/+; *UAS-λEgfr*/*dpp-LacZ* clones (bottom panels). In both types of clones there is precocious non-cell autonomous retinal differentiation (arrows) and ectopic *dpp* expression inside the clone.

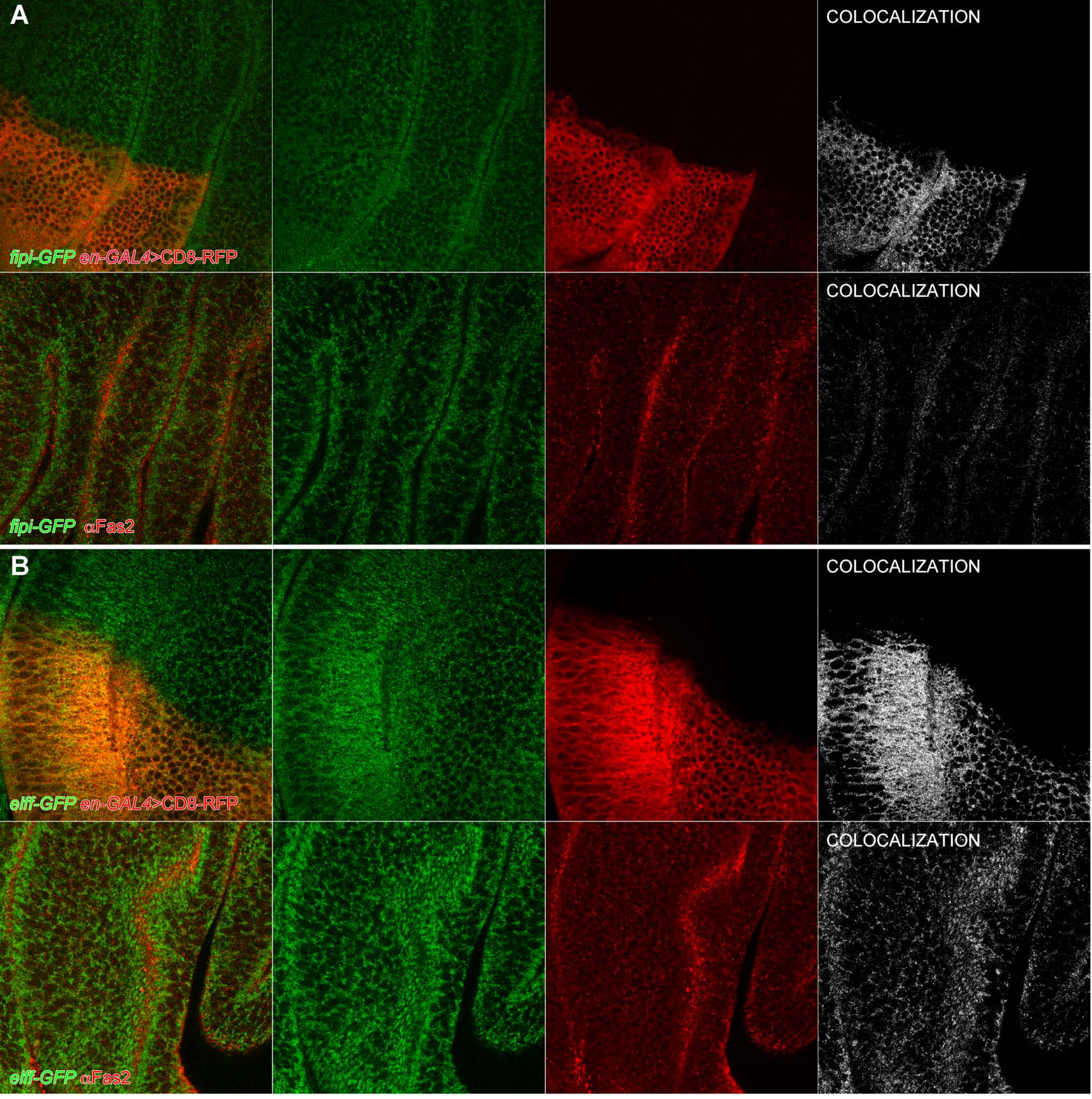
A
COLOCALIZATION
fipi-GFP en-GAL4>CD8-RFP
COLOCALIZATION
fipi-GFP αFas2
B
COLOCALIZATION
elff-GFP en-GAL4>CD8-RFP
COLOCALIZATION
elff-GFP αFas2

**S2 Fig. Fipi and Elff are expressed at the cell membrane and partially colocalize with Fas2 during late 3rd instar larva**

(A) Top: the protein trap line *fipi-EGFP-FLAG* (RRID:BDSC_60532) shows protein colocalization with the membrane marker CD8-RFP. 3rd instar wing imaginal disc stained with anti-Flag (green) and expressing *UAS-CD8-RFP* under the control of the *en-GAL4*/+ driver. ImageJ Colocalization and Colocalization Finder plugins, ratio: 50.0; threshold red: 100.0; threshold green: 100.0; Pearson's correlation: + 0,349. Bottom: the expression of *fipi-EGFP-FLAG* (RRID:BDSC_60532) (anti-Flag, green) partially colocalizes with Fas2 (Mab 1D4, red) in the wing imaginal disc during late 3rd instar larva. Expression of both proteins in different domains is better seen in the lateral views of cells at the imaginal disc folds. Note that the partial colocalization corresponds to the interface between Fas2 (red) and Fipi (green). ImageJ Colocalization and Colocalization Finder plugins, ratio: 50.0; threshold red: 100.0; threshold green: 100.0; Pearson's correlation: + 0,332.

(B) Top: the protein trap line *elff-EGFP-FLAG* (RRID:BDSC_60531) also shows colocalization with the membrane marker CD8-RFP. 3rd instar wing imaginal disc stained with anti-Flag (green) and expressing *UAS-CD8-RFP* under the control of the *en-GAL*/*+4* driver. ImageJ Colocalization and Colocalization Finder plugins, ratio: 50.0; threshold red: 100.0; threshold green: 100.0; Pearson's correlation: + 0,419. Bottom: *elff-EGFP-FLAG* (RRID:BDSC_60531) (anti-Flag, green) partially colocalizes with Fas2 (Mab 1D4, red) during late 3rd instar larva. The panel shows the cell profiles at folds near the prospective wing pouch. Note the colocalization at the apposition of red (Fas2) and green (Elff) signals. ImageJ Colocalization and

Colocalization Finder plugins, ratio: 50.0; threshold red: 100.0; threshold green: 100.0; Pearson's correlation: + 0,392.

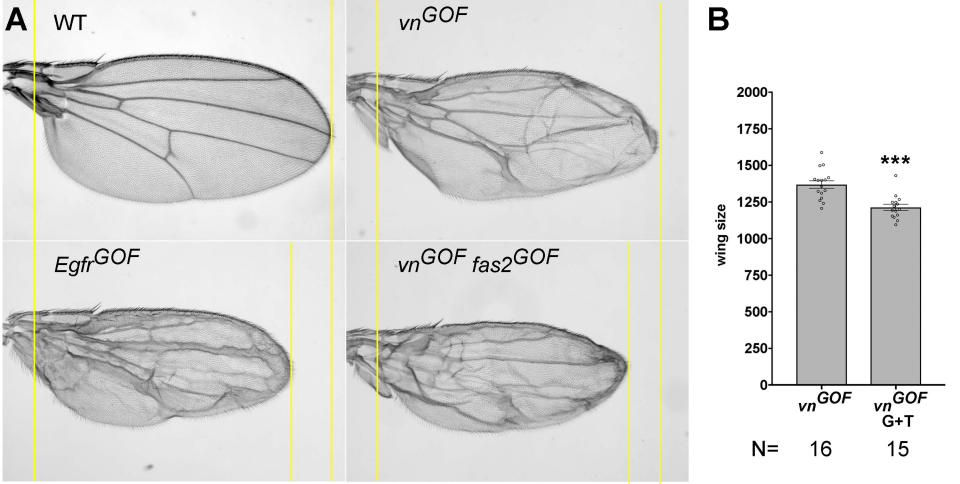

A
WT
vn^GOF
Egfr^GOF
vn^GOF fas2^GOF
B
wing size
2000
1750
1500
1250
1000
750
500
250
0
***
vn^GOF
vn^GOF G+T
N=
16
15

**S3 Fig. Over-expression of Fas2 enhances the EGFR activation caused by Vn over-expression**

(A) The gain of function for EGFR (*Egfr*$^{GOF}$, *MS1096-GAL4*/+*; Egfr*$^{\lambda top}$/+; bottom left) during imaginal wing disc growth produces adult wings smaller than normal (WT, *MS1096*/+, top left), and with a pronounced differentiation of extra-vein territory. A similar situation is attained by the over-expression of the EGF-like ligand Vein (*vn*$^{GOF}$; top right). Both, the reduction in wing size and the extra-vein phenotype caused by the over- expression of *UAS-vn* (under the control of the *MS1096-GAL4*/+ driver, *vn*$^{GOF}$) is enhanced by the simultaneous over-expression of *fas2* (*vn*$^{GOF}$ *fas2*$^{GOF}$, *MS1096*/+*; UAS-vn*/*UAS- fas2*$^{GPI}$ *UAS-fas2*$^{TRM}$; bottom right).

(B) Quantification of wing area in over-expression conditions for Vn (*vn*$^{GOF}$) and Vn plus Fas2$^{GPI}$ Fas2$^{TRM}$ (*vn*$^{GOF}$ G+T) under the control of the *MS1096-GAL4*/+ driver. All pictures heterozygous *MS1096-GAL4*/+ females raised at 25ºC. Wing size is area in µm$^{2}$/10$^{3}$.

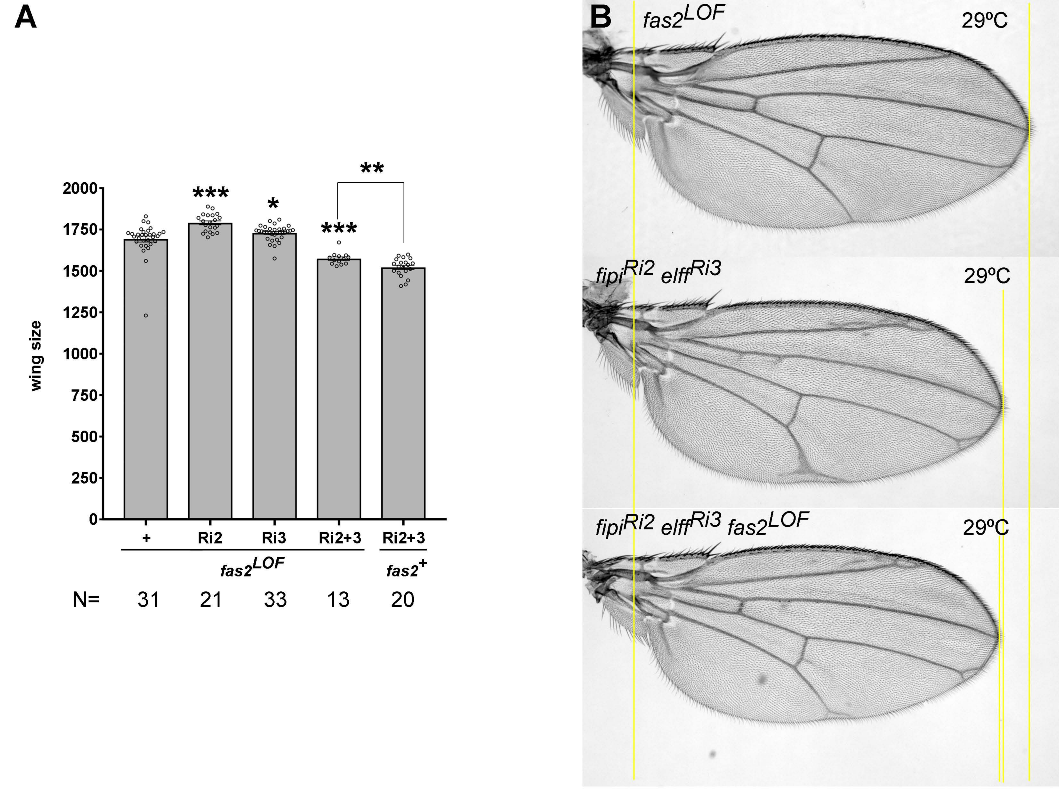
A
B
wing size
2000
1750
1500
1250
1000
750
500
250
0
***
*
***
**
+
Ri2
Ri3
Ri2+3
Ri2+3
fas2LOF
fas2+
N=
31
21
33
13
20
fas2LOF
29ºC
fipiRi2 elffRi3
29ºC
fipiRi2 elffRi3 fas2LOF
29ºC

**S4 Fig. *fipi* and *elff* LOF conditions are epistatic to the *fas2* LOF**

(A) Inhibition of *fipi* (*RNAi* RRID:BDSC_42589, Ri2) or *elff* (*RNAi* RRID:VDRC_32576, Ri3) in *MS1096-GAL4*/+; *UAS-fas2*$^{RNAi\#28990}$ females normalizes adult wing size. The double inhibition of *fipi* and *elff* expression in the *fas2* background (*fas2RNAi* RRID:BDSC_28990) shows a phenotype slightly suppressed. The combination *fipi elff* same as in Fig 5C. All individuals are heterozygous *MS1096-GAL4*/+ females raised at 25ºC. Wing size is area in $\mu m^2/10^3$.

(B) The inhibition of *fas2* by the expression of *RNAi* (RRID:BDSC_28990) in *MS1096-GAL4*/+ females (*fas2*$^{LOF}$) raised at 29ºC does not produce alterations in the wing vein pattern (with the exception of missing cross-veins in some individuals). Simultaneous inhibition of *fipi* and *elff* (*RNAi* RRID:BDSC_42589 and *RNAi* RRID:VDRC_32576) in *MS1096-GAL4*/+ females (*fipi*$^{Ri2}$ *elff*$^{Ri3}$) raised at 29ºC causes the differentiation of extra-vein tissue, while the simultaneous inhibition of *fas2 fipi* and *elff* expression (*fipi*$^{Ri2}$ *elff*$^{Ri3}$ *fas2*$^{LOF}$) reduces, but does not eliminate, the formation of extra-veins. Wing size is similar with or without *fas2* inhibition. All pictures heterozygous *MS1096-GAL4*/+ females raised at 29ºC.

Strains used for MARCM and coupled-MARCM analysis:

*y QS13F FRT19A/FM7-GFP; QF-ET40 QUAS-mtdTomato/CyO*

*y QS13F FRT19A; UAS-fas2*$^{GPI}$ *UAS-fas2*$^{TRM}$

*w hsp70-flp Tub-GAL80 FRT19A; Tub-GAL4 UAS-GFP/TM6B*

*y ey-flp Tub-GAL80 FRT19A; Tub-Gal4 UAS-GFP/TM6B*

Strains for FLP-OUT clone analysis:

*y ey-flp; Act5C-FRTy*$^{+}$*FRT-GAL4 UAS-GFP/CyO*

*y w hsp70-flp; Act5C-FRTy*$^{+}$*FRT-GAL4 UAS-GFP/CyO*

*y w hsp70-flp; Act5C-FRTy*$^{+}$*FRT-GAL4 UAS-GFP/CyO; Tub-miniCic::scarlet*

*y w hsp70-flp; Act5C-FRTy*$^{+}$*FRT-GAL4 UAS-GFP*; *puc-LacZ*$^{E69}$*/TM3*

*y w hsp70-flp; Act5C-FRTy*$^{+}$*FRT-GAL4 UAS-GFP; Act5C-FRT-polyA-FRT-LacZ.*

Strains used for time controlled *UAS*-mediated expression:

*w; TubGAL80*$^{ts20}$*/CyO; Tub-GAL4 UAS-GFP/TM6B*

*MS1096-GAL4/FM6; Tub-GAL80*$^{ts20}$*/CyO*

*w; Tub-GAL80*$^{ts20}$*/CyO; hsp-GAL4*
